# Jaagsiekte sheep retrovirus infection induces changes in microRNA expression in the ovine lung

**DOI:** 10.1101/2021.10.27.466207

**Authors:** Maria Contreras Garcia, Anna E. Karagianni, Deepali Vasoya, Siddharth Jayaraman, Yao-Tang Lin, Ann R. Wood, Mark P. Dagleish, Chris Cousens, Mick Watson, Finn E. Grey, David J. Griffiths

## Abstract

Ovine pulmonary adenocarcinoma (OPA) is an infectious neoplastic lung disease of sheep caused by jaagsiekte sheep retrovirus. OPA is an important veterinary problem and is also a valuable large animal model for human lung adenocarcinoma. JSRV infects type 2 alveolar epithelial cells in the lung and induces the growth of tumors, but little is known about the molecular events that lead to the activation of oncogenic pathways in infected cells. MicroRNAs (miRNAs) are small RNA molecules of approximately 22 nucleotides with important roles in regulating gene expression in eukaryotes and with well-established roles in cancer. Here we used small-RNA sequencing to investigate the changes in miRNA expression that occur in JSRV-infected ovine lung. After filtering out low abundance miRNAs, we identified expression of 405 miRNAs, 32 of which were differentially expressed in JSRV-infected lung compared to mock-inoculated control lung. Highly upregulated miRNAs included miR-182, miR-183, miR-96 and miR-135b, which have also been associated with oncogenic changes in human lung cancer. Network analysis of genes potentially targeted by the deregulated miRNAs identified their involvement in pathways known to be dysregulated in OPA. We found no evidence to support the existence of miRNAs encoded by JSRV. This study provides the first information on miRNA expression in OPA and identifies a number of targets for future studies into the role of these molecules in the pathogenesis of this unique veterinary model for human lung adenocarcinoma.

**IMPORTANCE:** Ovine pulmonary adenocarcinoma is a neoplastic lung disease of sheep caused by jaagsiekte sheep retrovirus (JSRV). OPA is a significant welfare and economic concern for sheep producers and is a valuable large animal model for human lung adenocarcinoma. MicroRNAs are small RNA molecules of approximately 22 nucleotides with important functions in regulating gene expression in eukaryotes and with well-established roles in cancer. In this study, we examined the changes in microRNA expression that occur in the lung in response to JSRV infection. We identified differential expression of a number of host-encoded microRNAs in infected tissue, including microRNAs with roles in human cancer. We found no evidence that JSRV encodes a microRNA. This study provides new insights on the cellular response to JSRV infection in the ovine lung, which will inform future studies into the pathogenesis of OPA in sheep and its use as a model for human lung adenocarcinoma.

## INTRODUCTION

Ovine pulmonary adenocarcinoma (OPA) is a fatal neoplastic lung disease of sheep caused by jaagsiekte sheep retrovirus (JSRV) (1, 2). OPA is present in most sheep-rearing countries and results in significant economic losses for sheep producers. Sheep with clinically overt OPA appear thin, lose condition and are dyspnoeic when exercised. In addition, many affected sheep produce excess fluid which accumulates in the pulmonary airways and may be discharged from the nostrils when the head is lowered, a pathognomonic sign of the disease. OPA is a serious animal welfare issue.

JSRV infection in sheep induces carcinogenesis of secretory epithelial cells in the distal lung, where proliferating type 2 alveolar epithelial cells (AEC2s) comprise the large majority of tumor cells (3–5). Lesions observed in lungs of clinical cases of OPA resemble those seen in human lepidic pulmonary adenocarcinoma, a rare form of lung cancer previously called bronchioloalveolar carcinoma (6–8). In both malignancies, tumors are typically multifocal and found in the peripheral lung. OPA has therefore been suggested as a suitable model to study early oncogenic events in human lung adenocarcinoma (8, 9).

JSRV is unusual among oncogenic retroviruses in that its envelope (*env*) gene encodes a dominant oncoprotein that is capable of inducing lung tumors when expressed in sheep and mice (10–12). JSRV Env also induces features of cellular transformation when overexpressed in a variety of cell lines, including morphological changes and activation of protein tyrosine kinase signaling pathways such as PI3K-Akt and Raf-MEK-MAPK (13–16). These pathways are also found to be activated in OPA tumor cells in sheep (17–19) and in JSRV Env-transformed cells in murine models (11, 20). These interesting biological features present OPA as a unique animal model for understanding molecular events in lung carcinogenesis. However, many aspects of the interactions between JSRV and ovine lung tissue have yet to be investigated.

MicroRNAs (miRNAs) are short, non-coding RNAs of approximately 22 nucleotides that regulate gene expression post-transcriptionally (21). miRNAs were first described in the nematode *Caenorhabditis elegans* (22) and have since been discovered in organisms throughout the plant and animal kingdoms, in viruses and in green algae (23). Many miRNAs exhibit high sequence conservation in related species and some, such as let-7 (24), have homologs in distant species. Binding of miRNAs with their mRNA targets typically occurs by complementarity of the seed sequence (nucleotides 2-8 of the miRNA) to the 3′ -untranslated regions of genes (25). The targets of miRNAs may control various biological processes such as developmental timing, cell proliferation, cell death and tissue differentiation (26). Abnormal function of these processes leads to aberrant cell proliferation and many studies have investigated the potential roles of miRNAs in cancer (reviewed in (27, 28)).

Several viruses encode miRNAs, including herpesviruses, retroviruses and polyomaviruses (29). Virus-encoded miRNAs have been shown to influence cellular pathways related to their replication and pathogenesis. For example, the deltaretrovirus bovine leukemia virus (BLV), encodes 10 mature miRNAs (30). One of these, BLV-miR-B4, has similarity with host miR-29, and is associated with B-cell proliferation and oncogenesis, suggesting that viral miRNAs may play an important role in BLV-mediated transformation. Other retroviruses have been shown to modulate expression of cellular miRNAs in ways that may also influence pathogenesis; for example, mouse mammary tumor virus (MMTV) infection increases expression of members of the oncogenic cluster miR-17-92, which is known to be upregulated in a variety of human cancers, including mammary carcinomas (31). Similarly, human T-cell lymphotropic virus type 1 (HTLV-1) infection upregulates miR-93 and miR-130b, which target the pro-apoptotic Tumor Protein 53-Induced Nuclear Protein 1) (32).

Here, we used small RNA-sequencing to investigate miRNA expression in ovine lung tissue in the early stages of JSRV infection using an experimental lamb infection model. We identified differential expression of a number of miRNAs in infected lung tissue compared to lung tissue from mock-inoculated lambs. The differential expression of the majority of those miRNAs tested was confirmed by RT-qPCR in natural cases of OPA, suggesting an association between OPA and the upregulation of these miRNAs. We found no evidence to support the existence of miRNAs encoded by JSRV. Network analysis of the deregulated miRNAs identified their involvement in pathways previously identified as dysregulated in OPA (33). Collectively, this study provides new information on the host response to JSRV infection that may have relevance for understanding the pathogenesis of OPA and expands the limited knowledge of sheep miRNAs currently available.

## RESULTS

### Experimentally-induced cases of OPA were used for small RNA-Sequencing

In order to identify changes in miRNA expression in JSRV-infected lung tissue, we initially examined tissues from experimentally-infected specified pathogen-free (SPF) lambs and age-matched mock-inoculated control lambs. The use of experimentally-infected SPF lambs minimizes the effect of potentially confounding factors such as the presence of other infections or disease stage on miRNA expression. The experimental cases of OPA used have been described previously in studies examining JSRV target cells in the lung (4) and changes in mRNA transcription that occur following JSRV infection (33). Briefly, four 6-day old SPF lambs were experimentally-infected with JSRV, whereas four age-matched control lambs received cell culture medium. Each infected lamb was euthanized when signs of respiratory distress appeared (66 to 85 days post-inoculation) along with an age and sex-matched control lamb. OPA tumor lesions were observed in hematoxylin and eosin-stained lung tissue sections from infected lambs, and this was confirmed by immunohistochemistry (IHC) for the JSRV Env surface (SU) protein (4, 33). Such lesions were not present in lung tissue from mock-inoculated lambs. As is typical for this experimental infection model, the lung tissue of JSRV-infected animals used for RNA extraction was heterogeneous and by histological appearance comprised up to 10% of OPA-affected tissue in a background of normal tissue (4, 33).

### Detection of differentially expressed miRNAs in JSRV-infected lung tissue

RNA was extracted from eight discrete sites of the lungs of each animal and pooled for library preparation and sequencing. Small RNA-Seq generated over 19 million reads per sample and, following adapter trimming, over 87% of reads from each sample passed the quality threshold (Table 1). Comparison of the distribution of sequence length of the small RNAs revealed a similar pattern between JSRV-infected and mock-inoculated samples (Fig. 1A). The observed distribution is comparable to that reported in other small RNA-sequencing studies (34–36). The sequencing reads were then aligned to various categories of RNA (Fig. 1B). Over 64% of the total reads from each lamb mapped to miRNAs from miRBase (23), with the majority of the remainder mapping to other classes of RNA. Notably, a small proportion of reads (less than 0.003% per sample) mapped to the JSRV genome, raising the possibility that JSRV might encode small RNAs.

**Figure 1.**
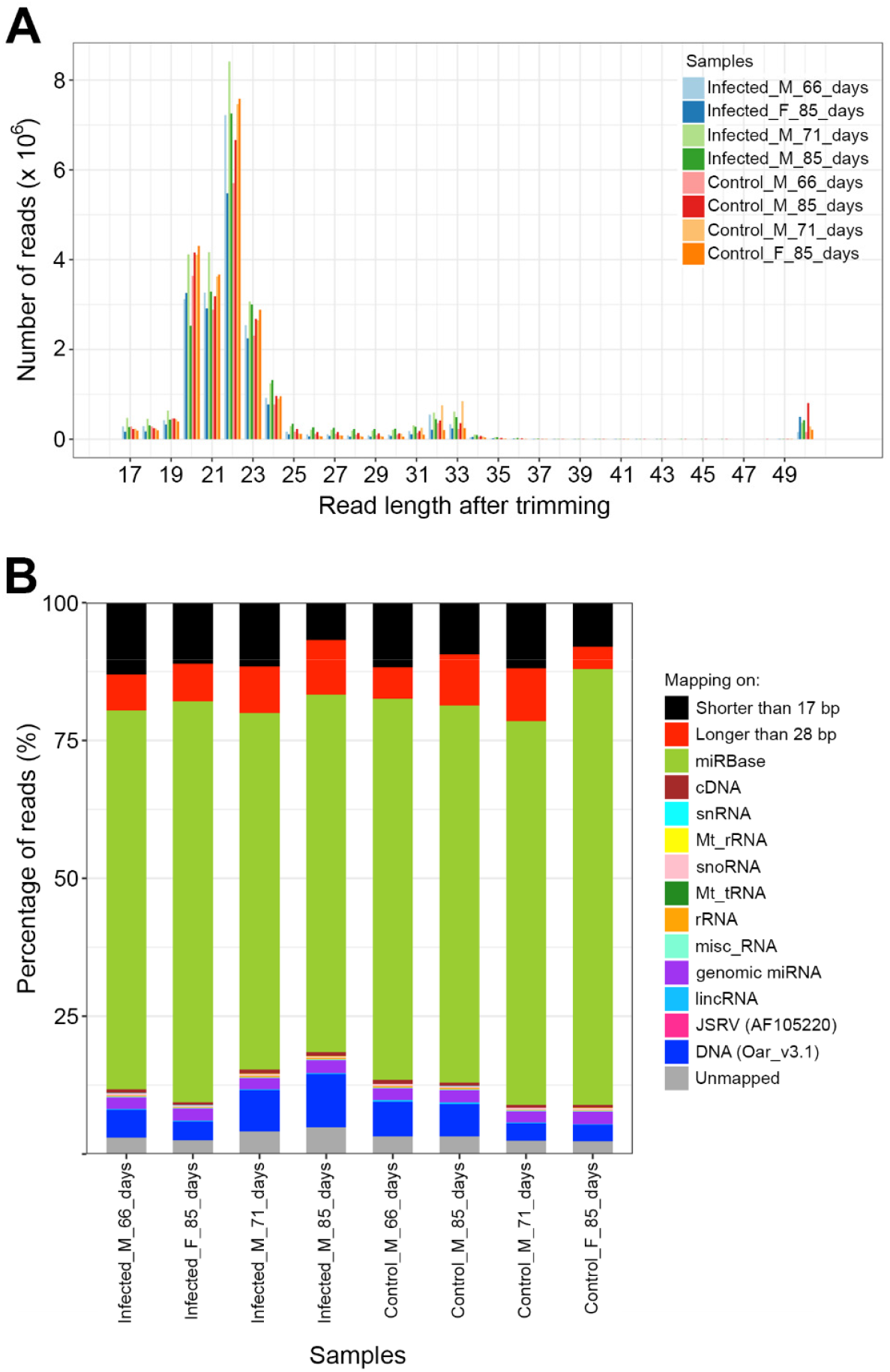
Statistics of small RNA sequencing. A, Read length distribution after quality trimming. The area with highest read abundance corresponds to RNAs with typical miRNA length (21 - 23 nt). B, Percentage of reads in each sample mapping to various RNA categories: snRNA, small nuclear RNA; Mt_rRNA, mitochondrial ribosomal RNA; snoRNA, small nucleolar RNA, Mt_tRNA, mitochondrial transfer RNA; rRNA, ribosomal RNA; misc_RNA, miscellaneous RNA; lincRNA, long intergenic non-coding RNA; JSRV, exogenous JSRV viral genome; DNA, sheep genome; Unmapped, reads not mapping to any of the previous categories. No statistically significant differences were observed between groups (t-test p<0.05).

**Table 1.** Read mapping statistics.

| Lamb <sup>a</sup> | Total No. of reads | No. of reads post quality trimming (%) | % total reads mapping to miRBase | % total reads mapping to JSRV genome |
| --- | --- | --- | --- | --- |
| Infected_M_66days | 23042889 | 18542545 (87.0) | 68.77 | 0.0022 |
| Infected_F_85days | 19063580 | 15645204 (88.9) | 72.68 | 0.0012 |
| Infected_M_71days | 29347531 | 23478540 (88.4) | 64.66 | 0.0028 |
| Infected_M_85days | 23400784 | 19495005 (93.2) | 64.86 | 0.0016 |
| Control_M_66days | 20380787 | 16831901(88.3) | 69.18 | 0.0011 |
| Control_M_85days | 23674153 | 19258905 (90.6) | 68.43 | 0.0008 |
| Control_M_71days | 25491448 | 20009358 (88.1) | 69.58 | 0.0010 |
| Control_F_85days | 23304071 | 20497050 (92.0) | 79.10 | 0.0008 |

<sup>a</sup> The lamb identifiers indicate the infection status, sex (male [M] or female [F]) and number of days post-inoculation when culled.

In total, 861 miRNAs were detected across all 8 lambs analyzed (Supplemental Data Set S1). We filtered out 456 miRNAs that were present at low abundance (mean normalized counts across samples lower than 50), as such reads are commonly thought to be unlikely to have biological significance. The remaining 405 miRNAs were used for further analysis. Of these, 318 miRNAs were not listed in the miRBase entry for *Ovis aries*, reflecting the relative paucity of coverage of sheep miRNAs in miRBase. Principal component analysis (PCA) was used to observe the variability among samples and to evaluate the clustering of samples as a first indication of differences between groups. JSRV-infected and mock-inoculated lambs clustered separately in the PCA plot (Fig. 2A) with greater variability within the JSRV-infected group. Notably, one JSRV-infected sample (Infected_F_85_days) was found to cluster more closely with the mock-inoculated group, indicating that the global miRNA expression pattern of this sample was more similar to mock-inoculated lambs. This finding is consistent with our previous analysis of mRNA transcription of the same samples (33), which found a lower level of JSRV infection in this lamb compared to the other lambs in this group.

**Figure 2.**
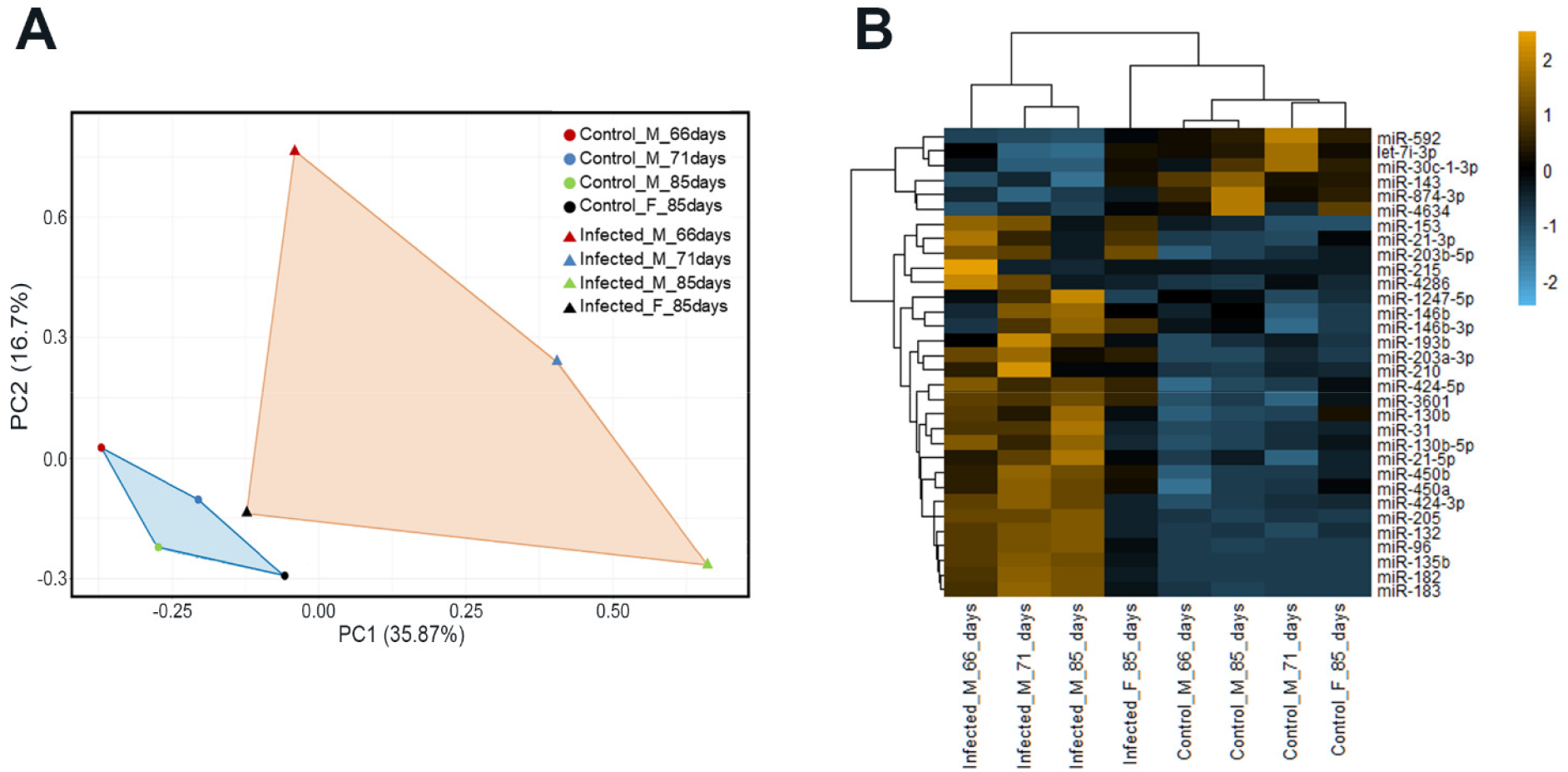
Differential expression of miRNAs in JSRV-infected compared to mock-inoculated lambs. A, Principal component analysis of miRNA expression in lung tissue (JSRV-infected lambs (n=4), mock-inoculated lambs (n=4)). Greater distance between samples in the plot indicates distinct expression patterns. Mock-inoculated samples formed a cluster towards the left of the plot, JSRV-infected samples covered a larger distance in both the x (PC1) and y axes (PC2). Sample Infected_F_85_days of the JSRV-infected group was closest to the mock-infected group in both PC1 and PC2, reflecting the lower level of infection in this lamb (33). The PCA plot indicates greater variability between JSRV-infected samples than among the mock-infected group and suggests global expression differences between the two groups. B, Heatmap of differentially expressed miRNAs (FDR<0.05, log_2_(fold change) ≥ 0.58 or ≤ -0.42) between lung tissue of JSRV-infected and mock-inoculated lambs. Dendrogram shows correlation clustering of individuals in groups. Legend represents values of log2 fold change.

Differential expression analysis was then performed to identify miRNAs with an altered expression pattern between JSRV-infected and mock-inoculated lambs. For this analysis we defined differentially expressed miRNAs to be those showing up- or down-regulation following JSRV infection with a false discovery rate (FDR) below 0.05, and a fold-change threshold established at ≤ 0.75 for downregulated miRNAs and ≥ 1.5 for upregulated miRNAs. The decision to establish these fold changes as significant was based on the low percentage of tumor tissue present in the samples to ensure that potential differences were detected while acknowledging the potential for false positives. Using these thresholds, we identified 32 miRNAs with significantly altered expression between JSRV-infected and mock-inoculated lambs (Fig. 2B; Table 2; Supplemental Data Set S1). Of these, 26 miRNAs were upregulated in JSRV-infected lambs and 6 were downregulated. As with the PCA, lamb Infected_F_85_days clustered more closely with the mock-inoculated lambs than with the other infected lambs, reflecting the lower level of infection in that animal.

**Table 2.** Differentially expressed miRNAs in JSRV-infected and mock-inoculated lung tissue.

| miRNA | Average counts<sup>1</sup> | log2(fold change)<sup>2</sup> | FDR<sup>3</sup> |
| --- | --- | --- | --- |
| miR-182<sup>4</sup> | 20880.16 | 3.20 | 0 |
| miR-183<sup>4</sup> | 7512.52 | 3.03 | 4.28E-84 |
| miR-96<sup>4</sup> | 343.43 | 3.15 | 5.94E-69 |
| miR-135b<sup>4</sup> | 70.94 | 4.82 | 1.28E-61 |
| miR-205<sup>4</sup> | 4802.59 | 1.29 | 1.01E-14 |
| miR-132 | 118.12 | 1.40 | 2.09E-11 |
| miR-450b | 4841.10 | 1.04 | 1.25E-08 |
| miR-424-3p | 188.42 | 1.07 | 4.56E-07 |
| miR-193b | 1564.57 | 0.92 | 5.68E-07 |
| miR-130b-5p | 130.98 | 1.13 | 7.09E-07 |
| miR-215 | 105.36 | 1.08 | 1.66E-06 |
| miR-31<sup>4</sup> | 252.35 | 0.91 | 2.06E-05 |
| miR-424-5p | 1250.31 | 0.87 | 2.15E-05 |
| miR-21-5p<sup>4</sup> | 28385.89 | 0.75 | 5.08E-05 |
| miR-1247-5p | 161.02 | 0.75 | 1.53E-04 |
| miR-450a | 1564.13 | 0.79 | 1.66E-04 |
| miR-203b-5p | 146.58 | 0.86 | 4.95E-04 |
| miR-3601 | 141.28 | 0.82 | 6.74E-04 |
| miR-210 | 547.69 | 0.70 | 8.33E-04 |
| miR-146b | 29159.79 | 0.61 | 8.61E-04 |
| miR-21-3p | 1407.30 | 0.69 | 1.22E-03 |
| miR-153 | 94.21 | 0.74 | 2.95E-03 |
| miR-130b | 690.02 | 0.70 | 2.98E-03 |
| miR-203a-3p | 963.90 | 0.61 | 6.78E-03 |
| miR-146b-3p | 58.90 | 0.72 | 1.08E-02 |
| miR-4286 | 90.54 | 0.61 | 4.02E-02 |
| miR-143 | 3626500.56 | -0.57 | 0 |
| miR-592 | 119.44 | -0.95 | 4.83E-05 |
| miR-30c-1-3p | 275.96 | -0.53 | 3.06E-02 |
| miR-874-3p | 514.64 | -0.57 | 3.13E-02 |
| let-7i-3p | 441.42 | -0.51 | 3.72E-02 |
| miR-4634 | 395.30 | -0.56 | 3.80E-02 |

<sup>1</sup>Mean normalized counts across all 8 samples. <sup>2</sup>Threshold established at log2(fold change) ≤ -0.42. <sup>3</sup> FDR
corrected by Benjamini-Hochberg, threshold FDR ≤ 0.05. <sup>4</sup>miRNAs also validated by RT-qPCR.

### Validation of RNA-Seq results by RT-qPCR

Nine miRNAs were selected for validation of differential expression by RT-qPCR. Seven of these (miR-21-5p, miR-31, miR-96, miR-135b, miR-182, miR-183 and miR-205) were identified based on the following criteria: fold change >1.5, mean normalized counts >50, coefficient of variation within groups <50%, FDR<0.01 and a minimum of five independent studies previously reporting their dysregulation or involvement in human lung cancer. In addition, miR-200b-5p and miR-503-5p were included in the validation panel. Although these two miRNAs did not meet the criteria for selection (miR-200b-5p: fold change of 1.46, FDR of 0.016 and mean normalized counts 274.65; miR-503-5p: fold change of 1.56, FDR of 0.0008 and mean normalized counts 11.72), they were of interest due to their known involvement, with validated targets, in lung cancer (37–39). miR-191 was chosen as the endogenous control for RT-qPCR due to its high expression level and low variance among all samples (mean normalized counts >150; coefficient of variance < 20%) and the lack of any reported involvement in lung cancer or viral infection. The stability of this miRNA was also validated by RT-qPCR and the results showed 5.6% coefficient of variance among the eight tested samples (data not shown).

Initially, we analyzed aliquots of the same RNA samples that had been used for small RNA sequencing. The results confirmed greater abundance of the nine selected miRNAs in the JSRV-infected group compared to the mock-inoculated control group (Fig. 3A, B) (p< 0.05). miR-183 had the greatest difference in expression between the two groups as it was detected in infected but not in control samples. The RT-qPCR analysis identified high variability within each group of lambs consistent with the results of small RNA-Seq.

**Figure 3.**
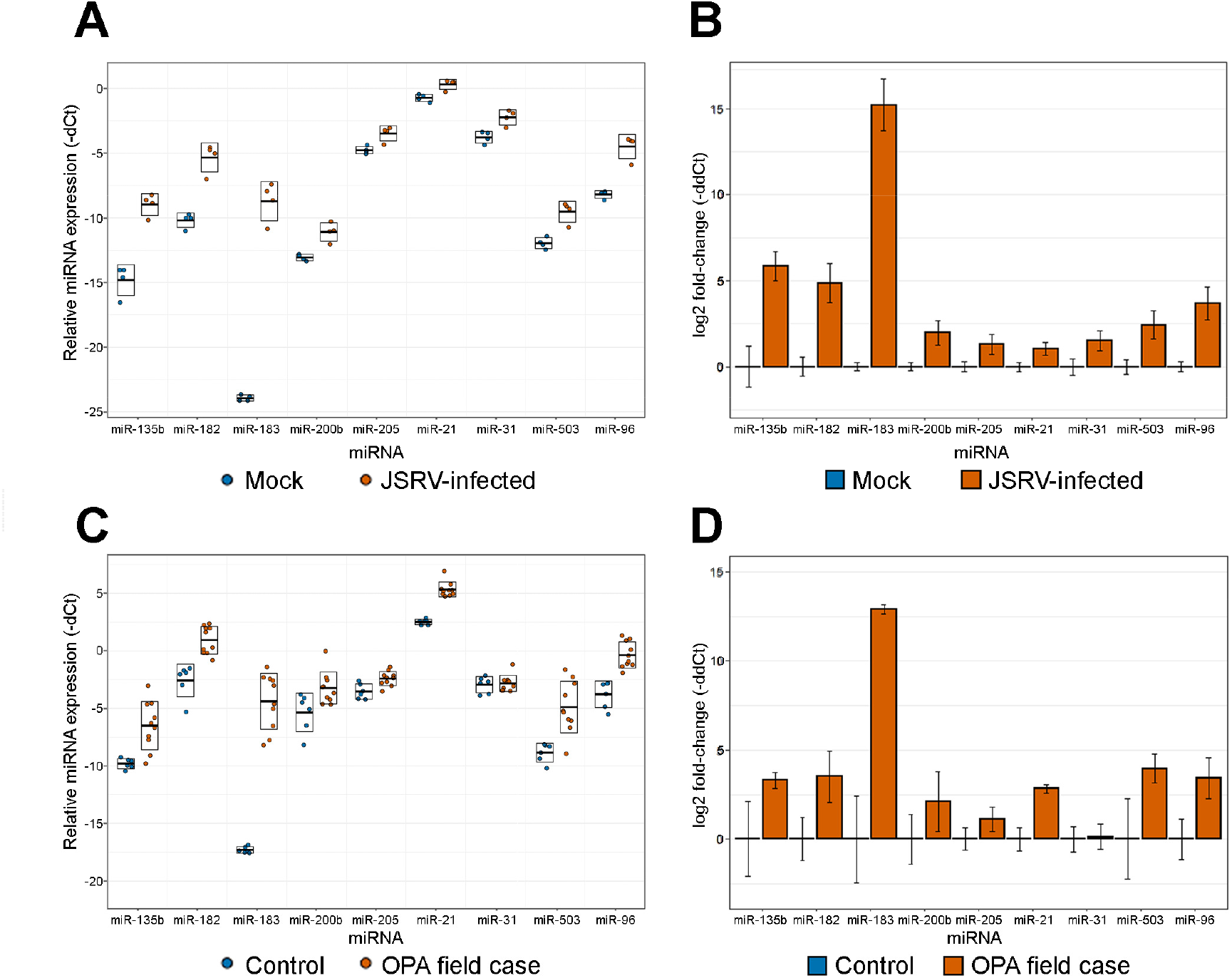
RT-qPCR analysis of miRNA expression in lung tissue of JSRV-infected and uninfected sheep. The expression of selected miRNAs was measured by RT-qPCR in lung tissue from JSRV-infected lambs (n=4) and mock-inoculated controls (n=4) (A, B), and in adult sheep with naturally acquired OPA (n=10) and clinically healthy control sheep (n=6) (C, D). A and C, Relative miRNA expression presented as -dCt (-Ct miR + Ct miR-191) of each individual sample per each miRNA assayed. Boxes display standard deviations of the mean, represented with a horizontal line. B and D, log_2_ fold-change between groups for each assayed miRNA, calculated following the ddCt method (94) and using miR-191 as an endogenous control. Error bars indicate standard deviation of miRNA expression within groups.

The tissues analyzed in the experimentally-infected lambs represent an early stage of OPA in young animals. In other diseases, the pattern of miRNA expression is known to vary with disease and developmental stage (40–42). Therefore, we next measured the expression of the nine selected miRNAs in lung tissues from adult sheep, including 10 naturally infected OPA cases and 6 clinically healthy sheep (Fig. 3C, D). All of the miRNAs in this panel were detected in both groups, with the exception of miR-183, which was not detected in samples from the clinically healthy control group. In addition, all but one of the miRNAs tested (miR-31) were significantly upregulated (p < 0.05) in OPA-affected sheep compared to healthy sheep. These findings indicate that miRNAs identified as upregulated in experimental cases of OPA are also upregulated in natural cases of OPA, increasing confidence that they are involved in the pathogenesis of OPA.

### JSRV does not encode a miRNA

Small RNA-sequencing reads were mapped to the genome of the JSRV isolate used in the experimental infections (JSRV_21_; GenBank Accession AF105220.1). The aligned reads were then visualized using Integrative Genomics viewer (IGV 2.3) in order to identify regions to which a disproportionately high number of reads aligned and that might therefore potentially encode viral miRNAs (Fig. 4).

**Figure 4.**
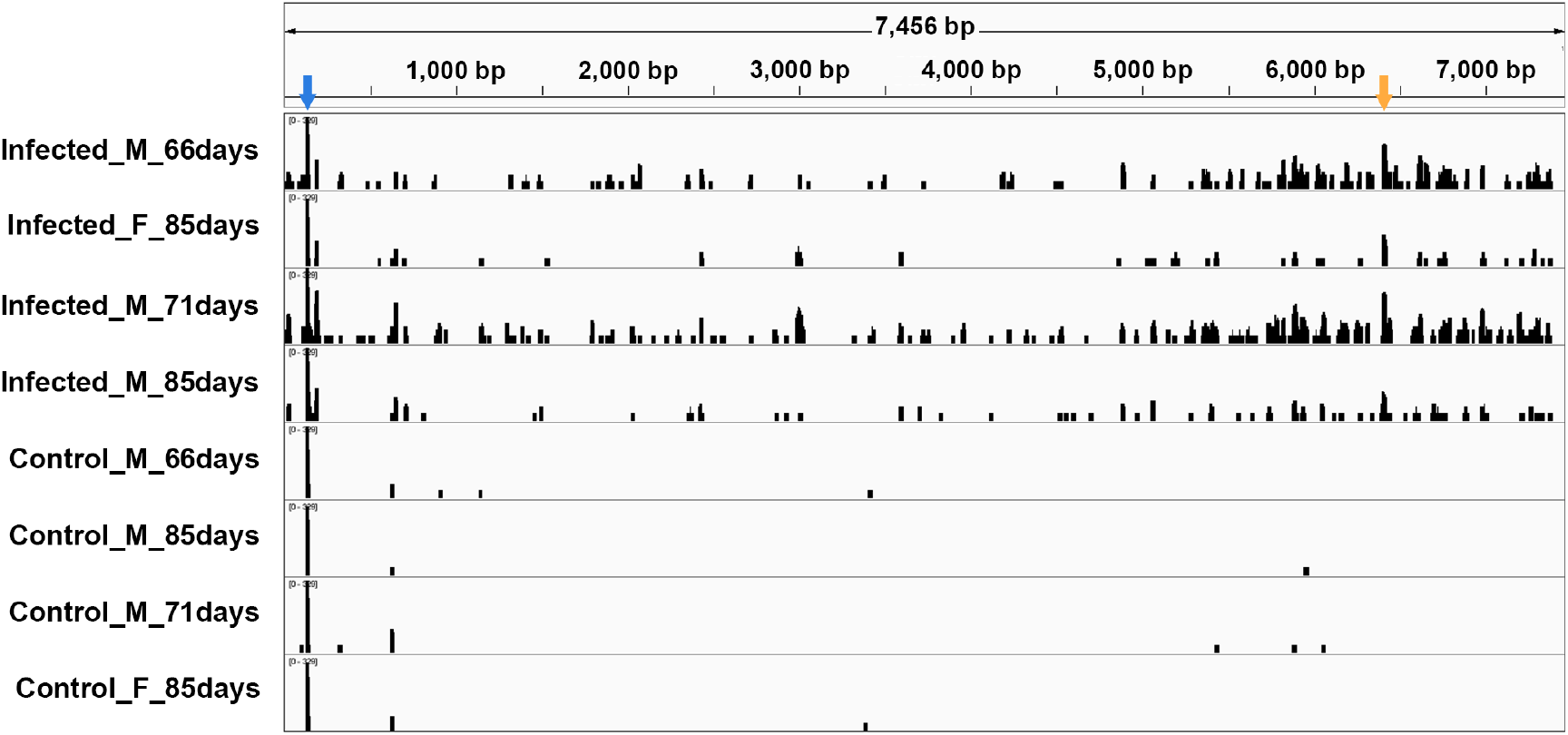
Small RNA-sequencing reads from JSRV-infected and uninfected lung tissue aligning to the JSRV genome sequence. The location of reads mapping to the JSRV genome in JSRV-infected (upper 4 panels) and mock-inoculated (lower 4 panels) sheep are shown. The orange arrow indicates the region around nucleotide 6396-6450 where a peak of reads was observed in JSRV-infected animals (see main text and Figure 5). The blue arrow indicates the mapping location of the tRNALys^1,2^ 3′-tRF molecule.

The most abundant read mapping to JSRV was CCCCACGUUGGGCGCCA, which was present in similar numbers in infected (166-329 counts) and uninfected (173-237 counts) animals. This read maps to a site immediately downstream of the JSRV 5′-LTR that is the binding site for a tRNA^Lys-1,2^ molecule that is used to prime reverse transcription (43, 44). Interestingly, this short RNA molecule represents the 3′-terminal 17 nt of the mature tRNA^Lys-1,2^, suggesting that it is a tRNA fragment (tRF). Recent studies have shown that tRFs are generated by specific cleavage from mature tRNAs and they may have several roles in regulating cellular gene expression (45).

Apart from the peak of reads representing the tRNA^Lys-1,2^ 3′-tRF, there were only 1-5 additional reads scattered across the genome in mock-inoculated animals. It appears likely that these originate from endogenous retroviruses related to JSRV (enJSRV), which are abundant in the sheep genome and transcribed in many tissues (46). These reads appear to map to JSRV in regions of high sequence similarity to enJSRV. In contrast, in JSRV-infected animals, reads were identified that mapped to several regions of the genome but in most cases at low coverage (Fig. 4). However, one exception to this was a region around nt 6400 of the JSRV genome, which accumulated 9-16% of the reads mapping to JSRV (10-49 reads per sample). The greater number of reads mapping to this region of the JSRV genome was not observed in mock-inoculated lambs, confirming that these reads were specific for JSRV-infected animals. The region around nt 6400 of the JSRV genome is part of the *env* gene, which is known to contain splice donor and acceptor sites for the expression of spliced transcripts (47). Potential structural and sequence features that could explain the higher number of reads in this area were investigated; however, no splice acceptors or other motifs were identified that could explain the greater abundance of reads mapping to this region.

The sequence surrounding nt 6400 of the JSRV genome was then evaluated for evidence that it might encode a miRNA. RNA folding prediction using RNAfold (48) identified the presence of a stem-loop structure of 54 nt in this region (Fig. 5A). Although pre-miRNA stem-loops are usually longer (∼ 70 nt), some retroviral miRNAs with shorter stem-loops have been reported (49). Nevertheless, by examining the stem-loop it can be observed that the potential miRNA sequence (UCAUACCAGGCUUCAGCUAUU) is not encoded entirely within the stem of the stem-loop structure, which is a requirement for DICER processing of miRNAs. This suggests that these reads are not derived from a genuine JSRV miRNA. To examine this potential miRNA experimentally, northern blotting was performed on RNA extracted from lung tissue samples from field cases of OPA-affected and clinically healthy sheep, and RNA from 293T cells transfected with plasmids that encode JSRV (pCMV2JS_21_ and pJSRV_21_ (43)) (Fig. 5). Hybridization was performed with three labeled probes: two to detect the -5p and -3p versions of the putative miRNA, and one to detect miR-191, which was used as a positive control.

**Figure 5.**
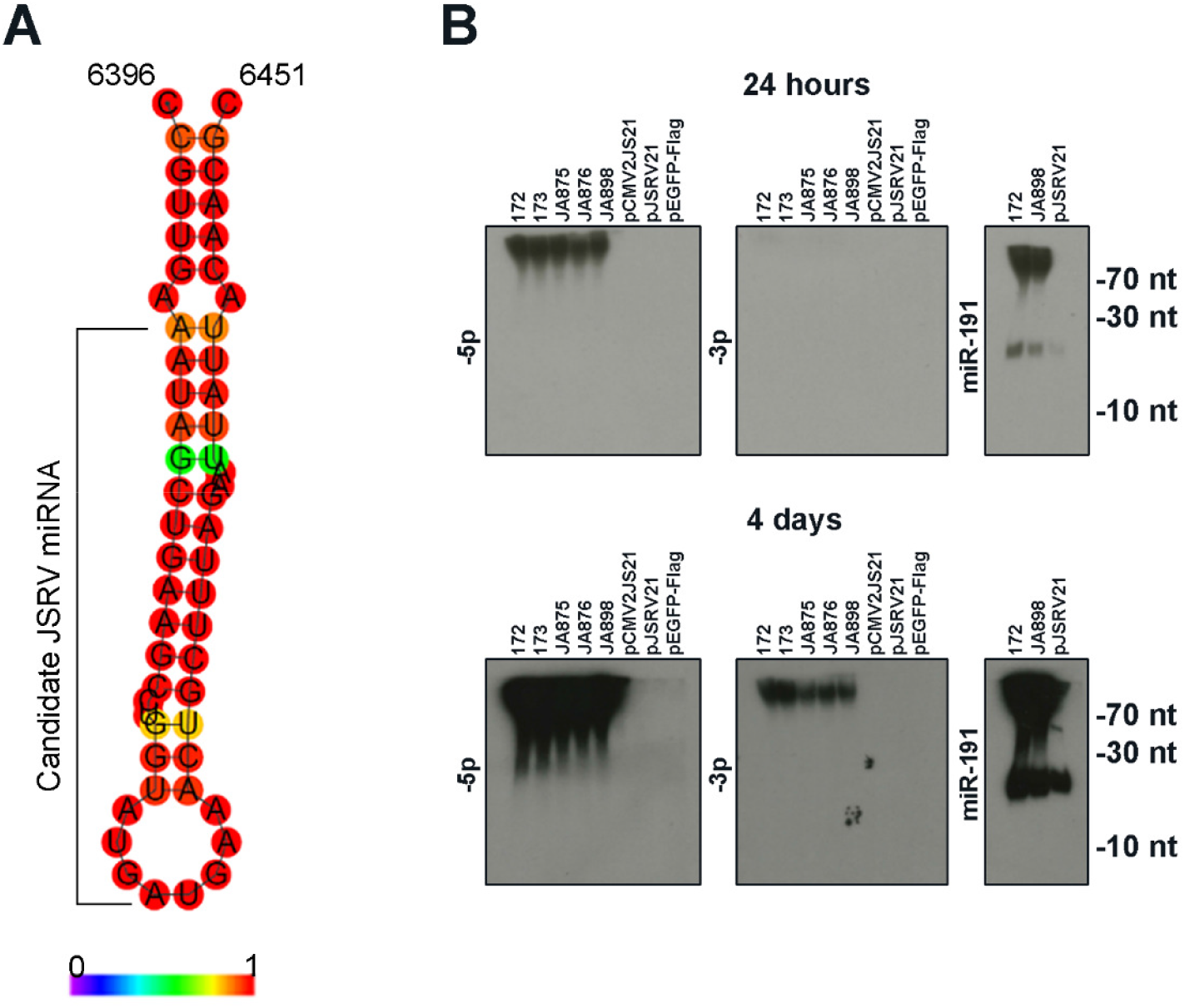
Evaluation of a potential miRNA encoded in the *env* region of the JSRV genome. A, Secondary structure prediction of the region of the JSRV_21_ genome (nucleotides 6396 – 6451) that encompasses the putative small RNA detected by small RNA sequencing. Colour legend shows the base-pair probabilities. The prediction was created with RNAfold (48). B, Northern blot analysis to detect the candidate JSRV miRNA. RNA from ovine lung tissue and transfected 293T cells was hybridized with probes for 5p and 3p arms of the putative JSRV miRNA and for cellular miR-191, as indicated. Samples 172 and 173 are control lung tissue from healthy sheep; samples JA875, JA876 and JA898 are OPA-affected lung tissue (from 3 independent field cases); pCMV2JS_21_ transfected 293T cells, pJSRV_21_ transfected 293T cells, pEGFP-Flag control transfected 293T cells. Upper panels show 24 hours exposure, lower panels show 4 day exposure. miR-191 was used as endogenous positive control.

The northern blot analysis revealed a band approximately 20 nt in size that hybridized with the labelled miR-191 probe (Fig. 5B), confirming successful detection of miR-191 and the integrity of the samples. In addition to the 20 nt band, a band of ∼ 70 nt in size was also detected in all samples. This band was observed more clearly after extended exposure of the blot (4-days) and may represent the pre-miRNA mir-191. In contrast, no bands of the expected miRNA size (21 nt) were observed when RNA samples were hybridized with probes for the -5p and -3p versions of the putative JSRV miRNA (Fig. 5B). Hybridization was performed with both -5p and -3p probes because detection of both forms of a miRNA would increase confidence in its existence. Both the -5p and -3p probes hybridized with a band at the top of lanes containing lung RNA from OPA-affected and control sheep, indicating that this sequence was not specific to JSRV-infected tissue. Given that the band was also not present in 293T samples, it is possible that the primer could also be binding to enJSRV sequences. Indeed, a BLASTN search found 18/21 nt identity between the probe and some enJSRV proviruses encoded in the sheep genome. Overall, the result of the northern blotting analysis did not provide any evidence to support the hypothesis that the sequencing reads mapping to the JSRV genome around nt 6400 are miRNAs. The potential origin of these sequencing reads and the reason for their apparent increased abundance remains unknown but may reflect an unidentified genomic feature of JSRV.

### Prediction and functional enrichment analysis of miRNA target genes

The small RNA-Seq analysis was performed on the same tissues used previously for mRNA transcriptome analysis (33) and we therefore sought to perform an integrated analysis of the two datasets. In order to identify potential target sites, we first attempted to extract the sequences of 3′-untranslated regions of ovine genes from Ensembl. However, only approximately 5500 genes were annotated, which is not sufficient for a robust analysis. As an alternative, we used Ingenuity Pathway Analysis (IPA) software for target gene prediction and functional analysis. IPA identified that there was targeting information available for 30 of the 32 differentially expressed miRNAs (the exceptions were miR-3601 and miR-146b-3p). IPA reported 667 mRNAs related to 267 pathways as targeted by the differentially expressed miRNAs. The results, presented in Supplemental Data Set S2, indicate those mRNA targets that have been experimentally validated and those which are predicted with high or moderate confidence. The main diseases and biofunctions related to these target genes are shown in Fig. 6, which indicates significant enrichment of cancer-related pathways.

**Figure 6.**
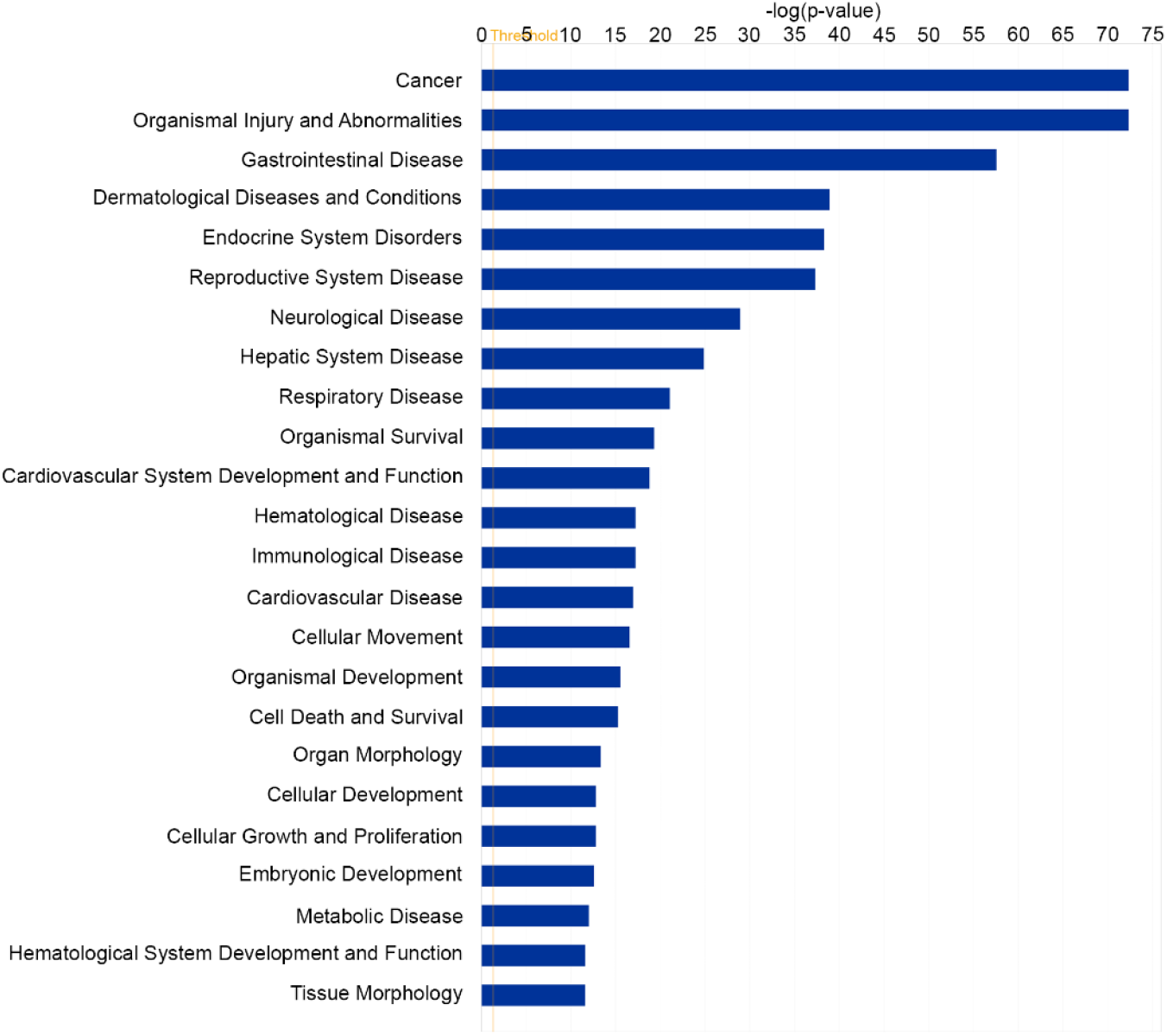
Diseases and Biofunctions associated with differentially expressed predicted target genes. The differentially expressed miRNAs were analyzed with Ingenuity Pathway Analysis software to identify potential target genes in the previously published dataset of genes differentially expressed in OPA (33). The figure shows the diseases and biofunctions associated with genes predicted to be targeted by miRNAs differentially expressed in JSRV-infected sheep lung, plotted by relative statistical significance. Significance values were calculated based on a right-tailed Fisher’s exact test, and the log(*P* value) is displayed on the horizontal axis of the bar chart. The taller the bar, the more significant the pathway effect.

## DISCUSSION

OPA is an important disease of sheep and a unique naturally-occurring model for human lung carcinogenesis. The involvement of miRNAs in cancer led us to investigate miRNA expression in OPA. We identified many miRNAs previously unidentified in sheep and 32 miRNAs that were differentially present in lung tissue from OPA-affected sheep compared to healthy sheep lung. In addition, our analysis found no evidence to support the existence of miRNAs encoded by JSRV.

405 miRNAs (with mean normalized abundance >50) were found to map to miRNAs in miRBase, of which 87 had been previously reported in sheep and 318 had been reported in other species (cattle, human and goat). There are relatively few annotated miRNAs in sheep due to the limited number of studies that have been performed to date. A study on miRNAs in ovine lentivirus infection (50) also used the approach of mapping RNA sequencing reads to miRBase and also found that most miRNAs mapped to miRNAs of related species, highlighting the known sequence conservation of miRNAs.

We filtered out miRNAs with a mean number of normalized counts (across all samples) lower than 50, in common with convention, because miRNAs with low counts are unlikely to have biological relevance and tend not to be validated in further RNA-Seq experiments or by RT-qPCR. Of the miRNAs filtered out in this way, when expression is compared between infected and control lambs only 9 had a Padj<0.05. Interestingly, most of these are related to other miRNAs we found to be differentially expressed. For example, miR-135a is related to miR-135b, miR-212 is from the same cluster as miR-132, miR-183-3p is the other arm of miR-183-5p, and miR-106a is from the miR-17 family. In addition, one low-abundance miRNA (miR-503-5p, mean abundance 11.72) was validated as upregulated in OPA by RT-qPCR.

Similarly, a small proportion of the sequencing reads (between 2 - 5%) did not map to annotated miRNAs or other classes of RNA. While it is possible that some of these unidentified reads may represent novel miRNAs, previous studies have suggested that such reads are typically of low abundance and that their detection in all samples is difficult (51–53). For that reason, we did not investigate those reads further.

Differential expression analysis of lung tissue samples successfully detected 32 differentially expressed miRNAs in JSRV-infected lambs compared to mock-inoculated controls. Of those miRNAs, only six were found to be downregulated with the remainder being upregulated in infected animals. While studies on miRNA dysregulation in cancer typically report a global downregulation of miRNAs in diseased tissue (54), the small number of specific downregulated miRNAs found here is most likely explained by the nature of the tissue samples used. Lung tissue samples from JSRV-infected lambs comprise small foci of tumor cells with the majority of the sample tissue, as much as 90%, being histologically normal. The high proportion of histologically normal tissue might therefore dilute the signal from downregulated miRNAs, rendering them undetectable in our analysis (55, 56). Such samples therefore also have greater sensitivity to detect upregulated miRNAs than downregulated miRNAs.

The heterogeneous nature of the tissue used for small RNA-Seq might also underlie some of the variability in miRNA expression in the JSRV-infected samples, as not all tissue samples will have the same proportion of tumor cells. Furthermore, in addition to tumor cells, lung tissue has many cell types, each of which might change in relative abundance within the affected tissue, and each of which might change their expression phenotype in the presence of JSRV-infected cells. Despite these complexities, the whole tissue approach provides a snapshot of global changes in miRNA expression in OPA-affected lung, and provides a starting point for the identification of cell type-specific changes using enriched or purified cell populations.

The target genes of the differentially expressed miRNAs have not been investigated in sheep, but some potential targets have been identified in humans. It may be difficult to extrapolate between human and sheep studies due to differences in gene sequences and in regulatory networks between the two species; however, given the transcriptional similarities between OPA and human non-small cell lung cancer (NSCLC) (33), comparison with human cancer studies can give clues to the roles of specific miRNAs. For example, miR-183, miR-182 and miR-96 were the most upregulated miRNAs in JSRV-infected lung tissue. In humans, these three miRNAs form a cluster, encoded on chromosome 7. In normal tissues, miR-183-182-96 cluster members have well-established roles in the development of sensory organs including the eye and ear (57, 58) and typically show low expression levels in other healthy tissues. However, these miRNAs consistently show increased expression in cancer and they are regarded as oncomiRs that are positively associated with cancer progression in a number of cancer types (59, 60). In human NSCLC, miR-183 has been reported to act to both promote and to restrict tumorigenesis (60–63).

In contrast to the established role of the miR-183-182-96 cluster in cancer, few studies have demonstrated its activation during infection, although roles have been reported in clonal expansion of helper T lymphocytes (64), suppression of natural killer cell function (65) and positive regulation of interferon responses (66). Notably, miR-183 is upregulated in tumors generated by infection with MMTV, another member of the betaretrovirus genus (31). Whether this is a result of the transformation process or the response to infection is unclear.

The miRNAs upregulated in OPA have targets in oncogenic pathways known to be activated in OPA. For example, the JSRV Env protein activates the PI3K-Akt-mTor signaling pathway (11, 13, 18, 67, 68) and miR-21 (69, 70), miR-183 (60) and miR-205 (71–73) all target *PTEN* and/or *PPP2CB* phosphatases, which negatively regulate this pathway. In addition, miR-31 targets *RASA1* and *SPRED1* in the MAPK pathway (74). Finally, miR-135b is known to regulate several factors involved in Hippo pathway signaling, including LATS2, NDR2, MOB1b and β-TrCP (75). Hippo signaling is crucial for alveolargenesis and dysregulation of this pathway has been described previously in OPA (33) and in human NSCLC (76). Collectively, the miRNAs found to be most differentially expressed in OPA are consistent with those that would be predicted based on previous work on human cancer and in murine models.

The enrichment analysis of the predicted target genes of the differentially expressed miRNAs was performed using IPA. This analysis revealed cancer-related functions to be highly enriched in the miRNA:mRNA data sets (Fig. 6; Supplemental Data S2). The IPA miRNA target gene analysis uses information from predicted targets of mammalian miRNAs, based on the observation that the majority of mammalian mRNAs are conserved targets of miRNAs (77–79). A similar analysis was reported previously with a dataset from cattle (80). Indeed, the evolvability of microRNA target sites among mammals (*i.e.*, the ‘proportion of evolutionarily changeable targets’) is estimated to be as low as 20% (79). However, as miRNA target sites are relatively uncharacterized in sheep, despite these results being consistent with previous studies on human lung cancer and OPA, further experimental validation is required to confirm the interactions.

Previous studies have identified miRNAs in other retroviruses, including BLV, HIV-1 and bovine and simian spumaretroviruses (30, 49, 81, 82). The small RNA-Seq data obtained in this study indicated increased abundance of reads mapping to a short region within the JSRV *env* gene, suggesting the possibility that JSRV might encode a miRNA. However, this was not supported by experimental analysis of sheep tissue or cell culture samples (Fig. 5), and we conclude that JSRV does not, in fact, encode a miRNA. This is consistent with small RNA-Seq studies of other betaretroviruses, including MMTV (31) and enzootic nasal tumor virus (83) and a bioinformatics screen of all retroviral genome sequences (30).

Although our primary focus in this study was to characterize miRNA expression in OPA, we also identified the presence of a 3′ -tRF derived from tRNA^Lys-1,2^ in sheep lung samples due to the presence of the complementary sequence in the primer binding site (PBS) region of the JSRV genome (43). A similar finding has been reported in an MMTV-infected mammary cell line (31). The potential for 3′ -tRFs to bind retroviral and retrotransposon genomes at the PBS is well established; for example, tRFs are able to repress expression of retrotransposons with a cognate PBS in preimplantation stem cells (84). It can be speculated that the tRF of tRNA^Lys-1,2^ could perhaps inhibit the replication of JSRV; for example, by competing for binding of the PBS with the mature full length tRNA. Similarly, this tRF might act to inhibit retrotransposition of enJSRV. However, it is unclear whether the tRF is sufficiently abundant in infected cells to exert such an effect. Further studies are necessary to directly address these questions. Notably, this tRF was present at a similar abundance in lung tissue from mock-inoculated and JSRV-infected lambs, suggesting that its generation was not stimulated by JSRV infection.

In summary, this study is the first to examine miRNA expression in OPA and has identified the differential expression of several miRNAs in affected lung tissue. These findings are a first step towards understanding the role of miRNAs and their predicted gene targets in OPA, which could be potentially exploited as biomarkers of the disease or as a tool to investigate the transformation process. In addition, this study contributes towards further characterization of host gene expression in OPA, which could aid the exploration of OPA as a human lung cancer model.

## MATERIALS AND METHODS

### In vivo studies

Samples used for RNA-Seq were available from a previous study in which SPF lambs were either experimentally-infected with JSRV_21_ (n=4) or mock-inoculated with culture supernatant (n=4) by intra-tracheal injection as described previously (4, 33). Lambs were born and housed under SPF conditions and were killed humanely by intravenous injection of sodium pentobarbital at the first clinical signs of respiratory distress. Age-matched controls were killed humanely at the same time. Post-mortem, tissue sections were collected from 24 different locations in the lungs, snap frozen in liquid nitrogen and stored at -80°C until analysis. Clinically healthy female adult sheep (n=6) and female OPA-affected adult sheep (n=10) were donated from local farms. Tissues from the lungs were collected at necropsy and stored at -80°C in the same way as samples from experimental cases. All procedures involving animals were performed with approval from the Moredun Research Institute Animal Welfare and Ethical Review Body and in conformance with the UK Animals (Scientific Procedures) Act 1986.

### RNA extraction

Lung tissue samples collected post-mortem and stored at -80°C were used to obtain RNA. From each experimental case, tissue sections from 8 different locations in the lungs were cut using a cryostat to represent the global state of the lung. Three 15 µm cryosections from each lung sample were collected in FastPrep lysing matrix D tubes (MP Biomedicals) and homogenized with a Precellys Evolution Tissue Homogenizer. RNA extraction was performed using the RNeasy Plus Micro procedure (Qiagen), according to the manufacturer’s protocol for isolation of total RNA including small RNAs. Samples derived from the same animal were pooled together and used for small-RNA sequencing. The quality and integrity of RNA samples was assessed using Nanodrop ONE and Agilent Bioanalyser (Agilent 2100) and only samples with RIN > 6.0, and 260/280 ratio > 1.9 were submitted for small-RNA sequencing.

### RNA sequencing and bioinformatics analysis

RNA-Seq was performed by Edinburgh Genomics (https://genomics.ed.ac.uk/). Library preparation was performed using a TruSeq Small RNA Sample Preparation kit (11 PCR cycles) and size selection of libraries was performed using Blue Pippin (Sage Science) selecting products of 120-163 bp. Libraries were quality checked by HS Qubit (Thermo) and Bioanalyzer (Agilent), before sequencing in a single lane of an Illumina Hiseq2500 (v4 High Output, 50-base single-end sequencing).

The quality of raw sequencing data was assessed, low quality reads were removed (Phred score < 28) and adapters were trimmed using Cutadapt (85). To select the optimum miRNA size, trimmed reads longer than 28 nt and shorter than 17 nt were filtered out. The selected reads were then mapped to ovine, bovine, human and caprine miRNA sequences of miRBase v.22.1 (23) using NovoAlign (Novocraft Technologies) with parameters: -m, -s 1, -t 30, -h 60. miRBase miRNA entries with identical sequences but different species of origin were grouped together in a single entry and the species prefix removed. In some instances, reads mapped to homolog miRNAs from different species which did not have an identical sequence. This mapping to highly similar but non-identical sequences is due to the presence of isomiRs, which are miRNA isoforms that originate from the same miRNA gene (21). In those instances, reads were not merged into a single entry. miRNA nomenclature was maintained so that it would reflect the miRBase entry name without the species prefix. The -3p and -5p suffixes were kept where present.

Bam files were then analyzed, and raw reads were normalized in MATLAB (MathWorks) based on size factors. This normalization approach consists of considering a size factor for each library to compute the effective library size. The size factors are calculated by taking the median of the ratios of observed counts to those of a reference sample, whose counts are determined by calculating the mean of each gene across all samples (86). By dividing the counts of each library by the corresponding size factors, all counts are in the same scale, making them comparable. A threshold was then established to remove reads with low numbers of normalized counts from the differential expression analysis. An average of 50 normalized counts across samples was established as a cut-off, based on published literature (87–90).

Differential expression analysis and statistics were also performed in MATLAB using the negative binomial model (nbintest) with the ‘Constant’ option. A threshold FDR < 0.05 was established as statistically significant. Data used for PCA plots consisted of normalized counts of all miRNAs above the established threshold of an average of 50 normalized counts. PCA plots were created using RStudio, using the pam function of ggplot2 (ggfortify) (91). Data used for heatmaps consisted of the differentially expressed miRNAs. Heatmaps were created using RStudio, using the pheatmap function. Default options of the pheatmap function were used, with the following exceptions: scale was set to “row”, column clustering distance was “correlation”, row clustering distance was “Euclidean”, clustering method was “average”.

Reads were also mapped to JSRV_21_ (GenBank accession AF105220.1) and enJSRV (EF680301.1). Reads were counted using a custom-made Perl Script that allowed only 1 mismatch. Alignments were visualized using Integrative Genomics Viewer (Broad Institute; https://software.broadinstitute.org/software/igv/).

### Reverse transcription-quantitative polymerase chain reaction (RT-qPCR) for detection of miRNAs

The expression of miRNAs was assessed using RT-qPCR with the TaqMan Advanced miRNA cDNA Synthesis Kit (Applied Biosystems) in conjunction with Taqman Advanced miRNA assays (Applied Biosystems, assays: hsa-miR-135b-5p, hsa-miR-182-5p, hsa-miR-183-5p, hsa-miR-200b-5p, hsa-miR-205-5p, rno-miR-21-5p, hsa-miR-31-5p, mmu-miR-503-5p, hsa-miR-96-5p, hsa-miR-191-5p). 10 ng of RNA were used per reaction. In brief, the protocol requires a poly-A tailing reaction and a ligation reaction to be performed before reverse transcription, allowing universal primers to be used in the reverse transcription reaction. A universal amplification reaction followed the reverse transcription, which increased the starting cDNA input for qPCR. All these reactions were performed according to manufacturer’s instructions using a Biometra T-one thermocycler (Analytik Jena). Quantitative PCRs (qPCRs) were performed using the Taqman universal PCR mastermix (Applied Biosystems) as instructed in the manufacturer’s protocol and each of the sample-target combinations were assayed in duplicate. qPCR reactions were run in the ABI 7500 Real time PCR system (Applied Biosystems) with cycling conditions: 2 minutes at 50°C, 10 minutes at 95°C, and 40 cycles at 95°C for 15 seconds followed by 1 minute at 60°C.

### Validation of RT-qPCR parameters

Efficiency (E) of qPCR was assessed by making six serial 5-fold dilutions of the template in molecular grade water and performing the qPCR reaction in duplicate for each dilution. Efficiency was then calculated by plotting the Ct values of each reaction against the logarithm of the template concentration and fitting a line through the points in the plot. The slope of the fitted line was used to calculate the efficiency percentage using the following equation: E = (10^(-1/slope)^)-1. The calculated efficiencies ranged from 92.35% to 107.71%.

### RNA-structure analysis

Prediction of the stem-loop structure in the putative miRNA encoded in the region of nt 6400 of the JSRV genome was performed using RNAfold (48) (http://rna.tbi.univie.ac.at/cgi-bin/RNAWebSuite/RNAfold.cgi).

### Northern blotting

RNA samples for northern blotting were concentrated by ethanol precipitation to achieve concentrations greater than 800 ng/µl. Northern blotting was performed as previously described (92). Briefly, electrophoresis was performed with a 15% acrylamide gel and gels were stained with ethidium bromide and analyzed under ultraviolet light to visualize tRNAs and assess sample degradation. RNA was transferred to Amersham Hybond-N+ membranes (GE Healthcare) with a mini-Protean blotting system (Biorad) in 0.5× Tris Borate buffer for 45 minutes at 30 V, followed by 15 minutes at 35 V and 15 minutes at 40 V. The marker bands corresponding to 70 nt (tRNAs), 30 nt (band produced by xylene cyanol) and 10 nt (band produced by bromophenol blue) were marked with pencil in the membranes as size indicators. Crosslinking of the RNA to the membranes was performed with a UV Stratalinker 1800 (Stratagene) at 1200 μJ/m^2^. Membranes were left to dry overnight at room temperature.

[γ-^32^P]dATP-radiolabeled oligonucleotide probes were prepared by mixing 1 µl of 10 pmol oligonucleotide, 1 µl 10× T4 polynucleotide kinase (PNK) buffer (NEB), 6 µl of water, 1 µl T4 PNK (NEB) and 1 µl ATP [γ-32P] 250 µCi (Perkin Elmer) and incubating at 37°C for 1 h. The probes used were: JS-5p candidate JSRV miRNA TCATACCAGGCTTCAGCTATT, JS-3p candidate JSRV miRNA AATAATTCTAAAGCAGTTTCA, and miR-191/oar-miR-191/hsa-miR191 AGCTGCTTTTGGGATTCCGTTG. The labelled probes were then diluted in 40 µl of water and separated from free [γ-^32^P]dATP by gel filtration (Illustra Microspin columns; GE Healthcare). Dried membranes were pre-hybridized in 8 ml of pre-warmed ExpressHyb buffer (Clontech) for 1 h at 55°C before adding filtered ^32^P-labeled probes and incubating at 38.5°C overnight (Biometra OV5; Analytik Jena). The membranes were then washed with 40 ml pre-warmed washing buffer (2 × SSC (20 × SSC stock: 175.3 g/l sodium chloride and 88.2 g/l sodium citrate in water), 0.1% SDS (Sigma Aldrich) in water). Four washes were performed (20 mins each; 38.5°C) after which membranes were put in contact with filter paper, pre-soaked in 2 × SSC, and heat-sealed in Seal-o-meal. Labelled bands were then exposed to Biomax MR film (Carestream) inside a Biomax MS screen (Carestream) at -80°C for four days. The film was developed with an SRX-101A film processor (Konica Minolta).

### Analysis of differentially expressed miRNA and mRNA targets and functional assessment

The miRNA target genes were obtained using the MicroRNA Target Filter tool of Qiagen’s Ingenuity Pathway Analysis (IPA), as described previously (80, 93). The analysis is based on four target prediction databases TargetScan, miRecords, Ingenuity Expert Findings and TarBase. Only the target genes that were present in the differentially expressed dataset published previously (33) were included in the analysis. Both mRNA-Seq and miRNA-Seq data derive from the same tissue samples. A threshold was applied such that only negatively correlated miRNA:mRNA pairs were included in the analysis. The functional enrichment of miRNA -mRNA target was also performed using IPA software to identify enriched canonical pathways and biological processes related to this dataset.

### Data availability

The raw RNA-Seq reads (fastq data) of each sample are present in the European Nucleotide Archive with the accession number PRJEB47862.

## ACKNOWLEDGEMENTS

We thank Edinburgh Genomics (https://genomics.ed.ac.uk/) for conducting the RNA-Seq experiments and Jeanie Finlayson for excellent technical assistance. We thank the staff of the Moredun Research Institute Bioservices Division for exceptional animal care and the farmers who support our work through the donation of OPA-affected sheep. This project was supported by the Moredun Foundation, the Scottish Government Rural and Environment Science and Analytical Services Division (RESAS) and the Biotechnology and Biological Sciences Research Council (grants BB/L009129/1 and BB/L008505/1), including the Institute Strategic Program, and national capability awards to The Roslin Institute (grants BB/P013740/1, BB/P013759/1, BB/P013732/1, BB/J004235/1, BB/J004243/1, BBS/E/D/20002173, and BBS/E/D/20002174).

## REFERENCES

1. Fan H. 2003. Jaagsiekte Sheep Retrovirus and Lung Cancer. Springer Berlin Heidelberg.

2. Griffiths DJ, Martineau HM, Cousens C. 2010. Pathology and pathogenesis of ovine pulmonary adenocarcinoma. J Comp Pathol 142:260–83.

3. De las Heras M, de Martino A, Borobia M, Ortin A, Alvarez R, Borderias L, Gimenez-Más J. 2014. Solitary tumours associated with Jaagsiekte retrovirus in sheep are heterogeneous and contain cells expressing markers identifying progenitor cells in lung repair. J Comp Pathol 150:138–147.

4. Martineau HM, Cousens C, Imlach S, Dagleish MP, Griffiths DJ. 2011. Jaagsiekte sheep retrovirus infects multiple cell types in the ovine lung. J Virol 85:3341–55.

5. Murgia C, Caporale M, Ceesay O, Di Francesco G, Ferri N, Varasano V, de las Heras M, Palmarini M. 2011. Lung adenocarcinoma originates from retrovirus infection of proliferating type 2 pneumocytes during pulmonary post-natal development or tissue repair. PLoS Pathog 7:e1002014.

6. Palmarini M, Fan H. 2001. Retrovirus-induced ovine pulmonary adenocarcinoma, an animal model for lung cancer. J Natl Cancer Inst 93:1603–1614.

7. Perk K, Hod I. 1982. Sheep lung carcinoma: an endemic analogue of a sporadic human neoplasm. J Natl Cancer Inst 69:747–9.

8. Youssef G, Wallace WA, Dagleish MP, Cousens C, Griffiths DJ. 2015. Ovine pulmonary adenocarcinoma: a large animal model for human lung cancer. ILAR J 56:99–115.

9. Mornex JF, Thivolet F, De las Heras M, Leroux C. 2003. Pathology of human bronchioloalveolar carcinoma and its relationship to the ovine disease. Curr Top Microbiol Immunol 275:225–48.

10. Caporale M, Cousens C, Centorame P, Pinoni C, De las Heras M, Palmarini M. 2006. Expression of the jaagsiekte sheep retrovirus envelope glycoprotein is sufficient to induce lung tumors in sheep. J Virol 80:8030–7.

11. Linnerth-Petrik NM, Santry LA, Yu DL, Wootton SK. 2012. Adeno-associated virus vector mediated expression of an oncogenic retroviral envelope protein induces lung adenocarcinomas in immunocompetent mice. PLoS One 7:e51400.

12. Wootton SK, Halbert CL, Miller AD. 2005. Sheep retrovirus structural protein induces lung tumours. Nature 434:904.

13. Maeda N, Fu W, Ortin A, de las Heras M, Fan H. 2005. Roles of the Ras-MEK-mitogen-activated protein kinase and phosphatidylinositol 3-kinase-Akt-mTOR pathways in Jaagsiekte sheep retrovirus-induced transformation of rodent fibroblast and epithelial cell lines. J Virol 79:4440–50.

14. Maeda N, Palmarini M, Murgia C, Fan H. 2001. Direct transformation of rodent fibroblasts by jaagsiekte sheep retrovirus DNA. Proc Natl Acad Sci U S A 98:4449–54.

15. Rai SK, Duh FM, Vigdorovich V, Danilkovitch-Miagkova A, Lerman MI, Miller AD. 2001. Candidate tumor suppressor HYAL2 is a glycosylphosphatidylinositol (GPI)-anchored cell-surface receptor for jaagsiekte sheep retrovirus, the envelope protein of which mediates oncogenic transformation. Proc Natl Acad Sci U S A 98:4443–8.

16. Zavala G, Pretto C, Chow YH, Jones L, Alberti A, Grego E, De las Heras M, Palmarini M. 2003. Relevance of Akt phosphorylation in cell transformation induced by Jaagsiekte sheep retrovirus. Virology 312:95–105.

17. Archer F, Jacquier E, Lyon M, Chastang J, Cottin V, Mornex JF, Leroux C. 2007. Alveolar type II cells isolated from pulmonary adenocarcinoma: a model for JSRV expression in vitro. Am J Respir Cell Mol Biol 36:534–40.

18. Cousens C, Alleaume C, Bijsmans E, Martineau HM, Finlayson J, Dagleish MP, Griffiths DJ. 2015. Jaagsiekte sheep retrovirus infection of lung slice cultures. Retrovirology 12:31.

19. De Las Heras M, Ortin A, Benito A, Summers C, Ferrer LM, Sharp JM. 2006. In-situ demonstration of mitogen-activated protein kinase Erk 1/2 signalling pathway in contagious respiratory tumours of sheep and goats. J Comp Pathol 135:1–10.

20. Linnerth-Petrik NM, Walsh SR, Bogner PN, Morrison C, Wootton SK. 2014. Jaagsiekte sheep retrovirus detected in human lung cancer tissue arrays. BMC Res Notes 7:160.

21. Bartel DP. 2018. Metazoan MicroRNAs. Cell 173:20–51.

22. Lee RC, Feinbaum RL, Ambros V. 1993. The C. elegans heterochronic gene lin-4 encodes small RNAs with antisense complementarity to lin-14. Cell 75:843–54.

23. Griffiths-Jones S, Saini HK, van Dongen S, Enright AJ. 2008. miRBase: tools for microRNA genomics. Nuc Acids Res 36:D154–8.

24. Pasquinelli AE, Reinhart BJ, Slack F, Martindale MQ, Kuroda MI, Maller B, Hayward DC, Ball EE, Degnan B, Muller P, Spring J, Srinivasan A, Fishman M, Finnerty J, Corbo J, Levine M, Leahy P, Davidson E, Ruvkun G. 2000. Conservation of the sequence and temporal expression of let-7 heterochronic regulatory RNA. Nature 408:86–9.

25. Bartel DP. 2009. MicroRNAs: target recognition and regulatory functions. Cell 136:215–33.

26. Pichler M, Calin GA. 2015. MicroRNAs in cancer: from developmental genes in worms to their clinical application in patients. Br J Cancer 113:569–73.

27. Lee YS, Dutta A. 2009. MicroRNAs in cancer. Annu Rev Pathol 4:199–227.

28. Lin PY, Yu SL, Yang PC. 2010. MicroRNA in lung cancer. Br J Cancer 103:1144–8.

29. Kincaid RP, Sullivan CS. 2012. Virus-encoded microRNAs: an overview and a look to the future. PLoS Pathog 8:e1003018.

30. Kincaid RP, Burke JM, Sullivan CS. 2012. RNA virus microRNA that mimics a B-cell oncomiR. Proc Natl Acad Sci U S A 109:3077–82.

31. Kincaid RP, Panicker NG, Lozano MM, Sullivan CS, Dudley JP, Mustafa F. 2018. MMTV does not encode viral microRNAs but alters the levels of cancer-associated host microRNAs. Virology 513:180–187.

32. Yeung ML, Yasunaga J, Bennasser Y, Dusetti N, Harris D, Ahmad N, Matsuoka M, Jeang KT. 2008. Roles for microRNAs, miR-93 and miR-130b, and tumor protein 53-induced nuclear protein 1 tumor suppressor in cell growth dysregulation by human T-cell lymphotrophic virus 1. Cancer Res 68:8976–85.

33. Karagianni AE, Vasoya D, Finlayson J, Martineau HM, Wood AR, Cousens C, Dagleish MP, Watson M, Griffiths DJ. 2019. Transcriptional Response of Ovine Lung to Infection with Jaagsiekte Sheep Retrovirus. J Virol 93.

34. Chen F, Xu XY, Zhang M, Chen C, Shao HT, Shi Y. 2019. Deep sequencing profiles MicroRNAs related to Aspergillus fumigatus infection in lung tissues of mice. Journal of Microbiology, Immunol Infect 52:90–99.

35. Li N, You X, Chen T, Mackowiak SD, Friedlander MR, Weigt M, Du H, Gogol-Doring A, Chang Z, Dieterich C, Hu Y, Chen W. 2013. Global profiling of miRNAs and the hairpin precursors: insights into miRNA processing and novel miRNA discovery. Nuc Acids Res 41:3619–34.

36. Peng X, Gralinski L, Ferris MT, Frieman MB, Thomas MJ, Proll S, Korth MJ, Tisoncik JR, Heise M, Luo S. 2011. Integrative deep sequencing of the mouse lung transcriptome reveals differential expression of diverse classes of small RNAs in response to respiratory virus infection. mBio 2:e00198–11.

37. Pacurari M, Addison JB, Bondalapati N, Wan YW, Luo D, Qian Y, Castranova V, Ivanov AV, Guo NL. 2013. The microRNA-200 family targets multiple non-small cell lung cancer prognostic markers in H1299 cells and BEAS-2B cells. Int J Oncol 43:548–60.

38. Chen D-Q, Pan B-Z, Huang J-Y, Zhang K, Cui S-Y, De W, Wang R, Chen L-B. 2014. HDAC 1/4-mediated silencing of microRNA-200b promotes chemoresistance in human lung adenocarcinoma cells. Oncotarget 5:3333.

39. Yang Y, Liu L, Zhang Y, Guan H, Wu J, Zhu X, Yuan J, Li M. 2014. MiR-503 targets PI3K p85 and IKK-beta and suppresses progression of non-small cell lung cancer. Int J Cancer 135:1531–42.

40. Landgraf P, Rusu M, Sheridan R, Sewer A, Iovino N, Aravin A, Pfeffer S, Rice A, Kamphorst AO, Landthaler M, Lin C, Socci ND, Hermida L, Fulci V, Chiaretti S, Foa R, Schliwka J, Fuchs U, Novosel A, Muller RU, Schermer B, Bissels U, Inman J, Phan Q, Chien M, Weir DB, Choksi R, De Vita G, Frezzetti D, Trompeter HI, Hornung V, Teng G, Hartmann G, Palkovits M, Di Lauro R, Wernet P, Macino G, Rogler CE, Nagle JW, Ju J, Papavasiliou FN, Benzing T, Lichter P, Tam W, Brownstein MJ, Bosio A, Borkhardt A, Russo JJ, Sander C, Zavolan M, et al. 2007. A mammalian microRNA expression atlas based on small RNA library sequencing. Cell 129:1401–14.

41. Mancuso R, Hernis A, Agostini S, Rovaris M, Caputo D, Clerici M. 2015. MicroRNA-572 expression in multiple sclerosis patients with different patterns of clinical progression. J Transl Med 13:148.

42. Zheng W, Zhao J, Tao Y, Guo M, Ya Z, Chen C, Qin N, Zheng J, Luo J, Xu L. 2018. MicroRNA-21: A promising biomarker for the prognosis and diagnosis of non-small cell lung cancer. Oncol Lett 16:2777–2782.

43. Palmarini M, Sharp JM, de las Heras M, Fan H. 1999. Jaagsiekte sheep retrovirus is necessary and sufficient to induce a contagious lung cancer in sheep. J Virol 73:6964–72.

44. York D, Vigne R, Verwoerd D, Querat G. 1992. Nucleotide sequence of the jaagsiekte retrovirus, an exogenous and endogenous type D and B retrovirus of sheep and goats. J Virol 66:4930–4939.

45. Magee R, Rigoutsos I. 2020. On the expanding roles of tRNA fragments in modulating cell behavior. Nuc Acids Res 48:9433–9448.

46. Palmarini M, Mura M, Spencer TE. 2004. Endogenous betaretroviruses of sheep: teaching new lessons in retroviral interference and adaptation. J Gen Virol 85:1–13.

47. Nitta T, Hofacre A, Hull S, Fan H. 2009. Identification and mutational analysis of a Rej response element in Jaagsiekte sheep retrovirus RNA. J Virol 83:12499–511.

48. Langdon WB, Petke J, Lorenz R. 2018. Evolving better RNAfold structure prediction. Lecture Notes in Computer Science, 10781: 220–236.

49. Kincaid RP, Chen Y, Cox JE, Rethwilm A, Sullivan CS. 2014. Noncanonical microRNA (miRNA) biogenesis gives rise to retroviral mimics of lymphoproliferative and immunosuppressive host miRNAs. mBio 5:e00074–14.

50. Bilbao-Arribas M, Abendaño N, Varela-Martínez E, Reina R, de Andrés D, Jugo BM. 2019. Expression analysis of lung miRNAs responding to ovine VM virus infection by RNA-seq. BMC Genomics 20:62–62.

51. Farrell D, Shaughnessy RG, Britton L, MacHugh DE, Markey B, Gordon SV. 2015. The Identification of Circulating MiRNA in Bovine Serum and Their Potential as Novel Biomarkers of Early Mycobacterium avium subsp paratuberculosis Infection. PLoS One 10:e0134310.

52. Meng F, Hackenberg M, Li Z, Yan J, Chen T. 2012. Discovery of novel microRNAs in rat kidney using next generation sequencing and microarray validation. PLoS One 7:e34394.

53. Wake C, Labadorf A, Dumitriu A, Hoss AG, Bregu J, Albrecht KH, DeStefano AL, Myers RH. 2016. Novel microRNA discovery using small RNA sequencing in post-mortem human brain. BMC Genomics 17:776.

54. Lu J, Getz G, Miska EA, Alvarez-Saavedra E, Lamb J, Peck D, Sweet-Cordero A, Ebert BL, Mak RH, Ferrando AA, Downing JR, Jacks T, Horvitz HR, Golub TR. 2005. MicroRNA expression profiles classify human cancers. Nature 435:834–8.

55. Kim HK, Kim J, Korolevich S, Choi IJ, Kim CH, Munroe DJ, Green JE. 2011. Distinctions in gastric cancer gene expression signatures derived from laser capture microdissection versus histologic macrodissection. BMC Med Genomics 4:48.

56. Klee EW, Erdogan S, Tillmans L, Kosari F, Sun Z, Wigle DA, Yang P, Aubry MC, Vasmatzis G. 2009. Impact of sample acquisition and linear amplification on gene expression profiling of lung adenocarcinoma: laser capture micro-dissection cell-sampling versus bulk tissue-sampling. BMC Med Genomics 2:13.

57. Geng R, Furness DN, Muraleedharan CK, Zhang J, Dabdoub A, Lin V, Xu S. 2018. The microRNA-183/96/182 Cluster is Essential for Stereociliary Bundle Formation and Function of Cochlear Sensory Hair Cells. Sci Rep 8:18022.

58. Lumayag S, Haldin CE, Corbett NJ, Wahlin KJ, Cowan C, Turturro S, Larsen PE, Kovacs B, Witmer PD, Valle D, Zack DJ, Nicholson DA, Xu S. 2013. Inactivation of the microRNA-183/96/182 cluster results in syndromic retinal degeneration. Proc Natl Acad Sci U S A 110:E507–16.

59. Dambal S, Shah M, Mihelich B, Nonn L. 2015. The microRNA-183 cluster: the family that plays together stays together. Nucleic Acids Res 43:7173–88.

60. Ma Y, Liang AJ, Fan Y-P, Huang Y-R, Zhao X-M, Sun Y, Chen X-F. 2016. Dysregulation and functional roles of miR-183-96-182 cluster in cancer cell proliferation, invasion and metastasis. Oncotarget 7:42805–42825.

61. Kundu ST, Byers LA, Peng DH, Roybal JD, Diao L, Wang J, Tong P, Creighton CJ, Gibbons DL. 2016. The miR-200 family and the miR-183∼96∼182 cluster target Foxf2 to inhibit invasion and metastasis in lung cancers. Oncogene 35:173–86.

62. Li Y, Zhang H, Li Y, Zhao C, Fan Y, Liu J, Li X, Liu H, Chen J. 2018. MiR-182 inhibits the epithelial to mesenchymal transition and metastasis of lung cancer cells by targeting the Met gene. Mol Carcinog 57:125–136.

63. Zhang L, Quan H, Wang S, Li X, Che X. 2015. MiR-183 promotes growth of non-small cell lung cancer cells through FoxO1 inhibition. Tumour Biol 36:8121–6.

64. Stittrich AB, Haftmann C, Sgouroudis E, Kuhl AA, Hegazy AN, Panse I, Riedel R, Flossdorf M, Dong J, Fuhrmann F, Heinz GA, Fang Z, Li N, Bissels U, Hatam F, Jahn A, Hammoud B, Matz M, Schulze FM, Baumgrass R, Bosio A, Mollenkopf HJ, Grun J, Thiel A, Chen W, Hofer T, Loddenkemper C, Lohning M, Chang HD, Rajewsky N, Radbruch A, Mashreghi MF. 2010. The microRNA miR-182 is induced by IL-2 and promotes clonal expansion of activated helper T lymphocytes. Nat Immunol 11:1057–62.

65. Donatelli SS, Zhou JM, Gilvary DL, Eksioglu EA, Chen X, Cress WD, Haura EB, Schabath MB, Coppola D, Wei S, Djeu JY. 2014. TGF-beta-inducible microRNA-183 silences tumor-associated natural killer cells. Proc Natl Acad Sci U S A 111:4203–8.

66. Singaravelu R, Ahmed N, Quan C, Srinivasan P, Ablenas CJ, Roy DG, Pezacki JP. 2019. A conserved miRNA-183 cluster regulates the innate antiviral response. J Biol Chem 294:19785–19794.

67. Liu SL, Lerman MI, Miller AD. 2003. Putative phosphatidylinositol 3-kinase (PI3K) binding motifs in ovine betaretrovirus Env proteins are not essential for rodent fibroblast transformation and PI3K/Akt activation. J Virol 77:7924–35.

68. Palmarini M, Maeda N, Murgia C, De-Fraja C, Hofacre A, Fan H. 2001. A phosphatidylinositol 3-kinase docking site in the cytoplasmic tail of the Jaagsiekte sheep retrovirus transmembrane protein is essential for envelope-induced transformation of NIH 3T3 cells. J Virol 75:11002–9.

69. Yu S, Lu Z, Liu C, Meng Y, Ma Y, Zhao W, Liu J, Yu J, Chen J. 2010. miRNA-96 suppresses KRAS and functions as a tumor suppressor gene in pancreatic cancer. Cancer Res 70:6015–25.

70. Zhang JG, Wang JJ, Zhao F, Liu Q, Jiang K, Yang GH. 2010. MicroRNA-21 (miR-21) represses tumor suppressor PTEN and promotes growth and invasion in non-small cell lung cancer (NSCLC). Clinica Chimica Acta 411:846–852.

71. Cai J, Fang L, Huang Y, Li R, Yuan J, Yang Y, Zhu X, Chen B, Wu J, Li M. 2013. miR-205 targets PTEN and PHLPP2 to augment AKT signaling and drive malignant phenotypes in non-small cell lung cancer. Cancer Res 73:5402–15.

72. Wu H, Mo YY. 2009. Targeting miR-205 in breast cancer. Expert Opinion on Therapeutic Targets 13:1439–1448.

73. Li C, Yin Y, Liu X, Xi X, Xue W, Qu Y. 2017. Non-small cell lung cancer associated microRNA expression signature: integrated bioinformatics analysis, validation and clinical significance. Oncotarget 8:24564–24578.

74. Edmonds MD, Boyd KL, Moyo T, Mitra R, Duszynski R, Arrate MP, Chen X, Zhao Z, Blackwell TS, Andl T, Eischen CM. 2016. MicroRNA-31 initiates lung tumorigenesis and promotes mutant KRAS-driven lung cancer. J Clin Invest 126:349–364.

75. Lin CW, Chang YL, Chang YC, Lin JC, Chen CC, Pan SH, Wu CT, Chen HY, Yang SC, Hong TM, Yang PC. 2013. MicroRNA-135b promotes lung cancer metastasis by regulating multiple targets in the Hippo pathway and LZTS1. Nat Commun 4:1877.

76. Yeung B, Yu J, Yang X. 2016. Roles of the Hippo pathway in lung development and tumorigenesis. Int J Cancer 138:533–9.

77. Friedman RC, Farh KK, Burge CB, Bartel DP. 2009. Most mammalian mRNAs are conserved targets of microRNAs. Genome Res 19:92–105.

78. Naeem A, Zhong K, Moisá SJ, Drackley JK, Moyes KM, Loor JJ. 2012. Bioinformatics analysis of microRNA and putative target genes in bovine mammary tissue infected with Streptococcus uberis. J Dairy Sci 95:6397–408.

79. Xu J, Zhang R, Shen Y, Liu G, Lu X, Wu CI. 2013. The evolution of evolvability in microRNA target sites in vertebrates. Genome Res 23:1810–6.

80. Oliveira GB, Regitano LCA, Cesar ASM, Reecy JM, Degaki KY, Poleti MD, Felício AM, Koltes JE, Coutinho LL. 2018. Integrative analysis of microRNAs and mRNAs revealed regulation of composition and metabolism in Nelore cattle. BMC Genomics 19:126.

81. Ouellet DL, Plante I, Landry P, Barat C, Janelle ME, Flamand L, Tremblay MJ, Provost P. 2008. Identification of functional microRNAs released through asymmetrical processing of HIV-1 TAR element. Nuc Acids Res 36:2353–65.

82. Whisnant AW, Kehl T, Bao Q, Materniak M, Kuzmak J, Lochelt M, Cullen BR. 2014. Identification of novel, highly expressed retroviral microRNAs in cells infected by bovine foamy virus. J Virol 88:4679–86.

83. Wang B, Ye N, Cao SJ, Wen XT, Huang Y, Yan QG. 2016. Identification of novel and differentially expressed MicroRNAs in goat enzootic nasal adenocarcinoma. BMC Genomics 17:896.

84. Schorn AJ, Gutbrod MJ, LeBlanc C, Martienssen R. 2017. LTR-Retrotransposon Control by tRNA-Derived Small RNAs. Cell 170:61–71 e11.

85. Martin M. 2011. Cutadapt removes adapter sequences from high-throughput sequencing reads. EMBnetjournal 17:10–12.

86. Anders S, Huber W. 2010. Differential expression analysis for sequence count data. Genome Biology 11:R106.

87. Spornraft M, Kirchner B, Haase B, Benes V, Pfaffl MW, Riedmaier I. 2014. Optimization of extraction of circulating RNAs from plasma–enabling small RNA sequencing. PLoS One 9:e107259.

88. Motameny S, Wolters S, Nürnberg P, Schumacher B. 2010. Next generation sequencing of miRNAs–strategies, resources and methods. Genes 1:70–84.

89. Taxis TM, Bauermann FV, Ridpath JF, Casas E. 2017. Circulating microRNAs in serum from cattle challenged with bovine viral diarrhea virus. Frontiers in Genetics 8:91.

90. Koh W, Sheng CT, Tan B, Lee QY, Kuznetsov V, Kiang LS, Tanavde V. 2010. Analysis of deep sequencing microRNA expression profile from human embryonic stem cells derived mesenchymal stem cells reveals possible role of let-7 microRNA family in downstream targeting of hepatic nuclear factor 4 alpha. BMC Genomics 11:S6.

91. Clark EL, Bush SJ, McCulloch MEB, Farquhar IL, Young R, Lefevre L, Pridans C, Tsang HG, Wu C, Afrasiabi C, Watson M, Whitelaw CB, Freeman TC, Summers KM, Archibald AL, Hume DA. 2017. A high resolution atlas of gene expression in the domestic sheep (Ovis aries). PLoS Genet 13:e1006997.

92. McClure LV, Lin YT, Sullivan CS. 2011. Detection of viral microRNAs by Northern blot analysis. Methods Mol Biol 721:153–71.

93. Kramer A, Green J, Pollard J, Jr., Tugendreich S. 2014. Causal analysis approaches in Ingenuity Pathway Analysis. Bioinformatics 30:523–30.

94. Livak KJ, Schmittgen TD. 2001. Analysis of relative gene expression data using real-time quantitative PCR and the 2(-Delta Delta C(T)) Method. Methods 25:402–8.

